# Chemical compositions of essential oils, antimicrobial effect and antioxidant activity studies of *Hyoscyamus niger* L. from Turkey

**DOI:** 10.1101/2022.08.07.503024

**Authors:** Şule İnci, Pelin Yilmaz Sancar, Azize Demirpolat, Sevda Kirbag, Semsettin Civelek

## Abstract

In this study, essential oil components of the *Hyoscyamus niger* L. plant and their antimicrobial and antioxidant properties were determined. The essential oils were obtained separately from both the aerial parts and seeds of the plant using the hydrodistillation method. Antimicrobial activity was determined using the disk diffusion method. Total antioxidant level (TAS), total oxidant level (TOS), and 2.2-diphenyl-1-picrylhydrazyl (DPPH) radical scavenging capacity were detected for the antioxidant activity of the plant. The main essential oil components of the above-ground parts *Hyoscyamus niger* were determined as major components the phytol 52.09%, hexahydrofarnesyl acetone 19.66 %. *H. niger* seeds were hydrodistilled separately, obtaining yields of 0.7% (v w^-1^) of yellow oils. In the essential oils of *H. niger* seeds, forty-one components were identified representing 99.0% of the oils. According to the results of the analysis hexahydrofarnesyl acetone 46.36%, hexanal 9.05% as major components. It was determined that the methanol extracts of the above-ground *H. niger* inhibited the growth of pathogenic microorganisms at different rates (13-32 mm). TAS value of methanol extracts of *H. niger* was calculated as 3.7705 mmol, while the TOS value was calculated as 6.9403 μmol. It was determined that the scavenging effects of the DPPH radical increase, depend on increasing concentrations.

## INTRODUCTION

Today, medicinal plants, which have an important place in modern medicine and pharmacy, maintain their importance as a natural source of pharmaceutical raw materials due to the essential oils and secondary metabolites they contain (Akbaş et al., 2020).

The Solanaceae family, which includes many important medicinal plants, is a plant family with a wide variety of 98 genera and approximately 2700 species. Plants belonging to the Solanaceae family have been used by humans for many years as food, ornamental plants, and medicine (Guha and Maheshwari, 1966). The Solanaceae family, which contains many valuable plants rich in alkaloids with strong effects such as scopolamine, atropine, and hyoscyamine, therefore has great importance both medically and economically (Maiti et al., 2002).

*Hyoscyamus niger* (black henbane) is an annual or biennial herbaceous plant belonging to the Solanaceae family and is commonly found in rocky areas, uncultivated lands, and especially on roadsides (Orbak et al., 1998; Li et al., 2011; Yücel and Yılmaz, 2014). Hyoscyamus *niger* is an important resource for the pharmaceutical industry as it contains some important alkaloids, which are among the oldest drugs used in medicine (Orbak et al., 1998; Pudersell et al., 2012; Dehghan et al., 2012).

This plant is generally used in the treatment of stomach pains, ulcers, kidney and liver disorders, as well as it’s analgesic and antispasmodic effects (Tanker et al., 1998; John et al., 2010). However, with the consumption of the roots and leaves of the *Hyoscyamus niger*, some secondary metabolites in the plant can also cause poisoning by paralyzing the nerve endings of the parasympathetic system in humans (Orbak et al., 1998).

*H. niger* has been used as a hallucinogen since ancient times. Although it came to Europe in the middle ages, the hallucinogenic properties of this plant are even found in Ancient Greek writings. In particular, the seeds and leaves of the plant contain hyoscyamine, scopolamine, and other tropane alkaloids. Apart from the hallucinogenic properties of these chemicals, they are also responsible for the effects of ataxia and hypertension (Ugur, 2013). *H. niger* is an extremely poisonous plant and its consumption is undesirable. Because of this feature, it has also been used as a poison in history (Haas, 1995). Both placebo and *H. niger* methanolic extract to a group of 50 people in a clinic in Iran. It has been stated that the mixture applied to each patient three times a day for 6 days has beneficial effects in improving the symptoms of COVID-19. They defined that the clinical symptoms of COVID-19 such as dry cough, shortness of breath, sore throat, chest pain, fever, dizziness, headache, abdominal pain, and diarrhea were reduced with propolis plus *Hyoscyamus niger* L. extract than the placebo group (Kosari et al., 2021).

This study aims to determine the antimicrobial and antioxidant activity and essential oil composition of *Hyoscyamus niger*, a medicinal plant. In this study, while determining the essential oils in the plant, it will also be tried to determine the presence and limits of the antimicrobial effect on some bacterial and fungal species. Thus, it is aimed to contribute to the studies that have been done and will be done about providing raw materials for new therapeutics from plants that naturally grow in our country and have medicinal properties.

## MATERIALS AND METHODS

### Collection of the plant material

The plant material used in the study was collected from the Baskil district of Elazig province. Detailed locality information of the region where the plant was collected is given in Table 1. It was collected from its natural habitat in its intense flowering period in May-June, which is the flowering period of the plant, and dried in the shade. The specimens of the plant are preserved in the Herbarium (FUH) of the Faculty of Science, Firat University.

**Table 1.**
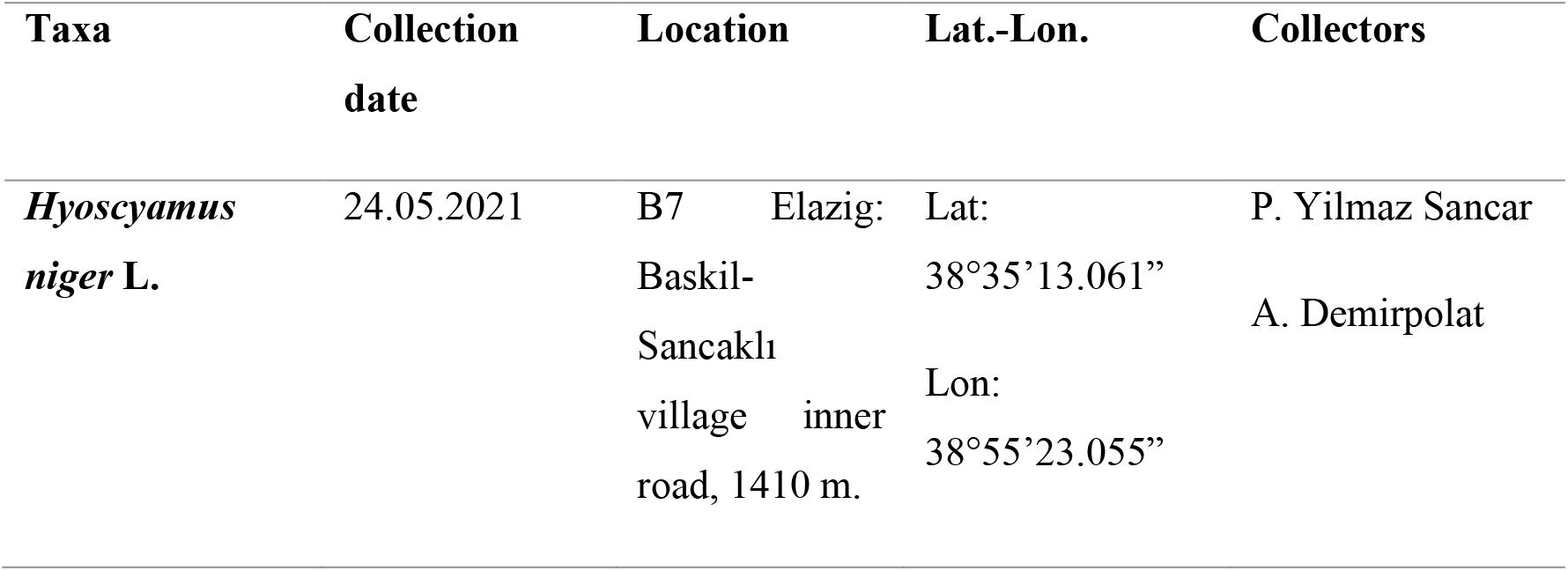
Detailed location information

### Isolation of the essential oils

In this study, the plants were air-dried. In recent years, effective methods have emerged in drying methods by drying in isothermal and non-isothermal systems (Demirpolat et al., 2021). The Hydrodistillation method was used to obtain the oil of the plant. Essential oil using a modified Clevenger-type apparatus for 4 h to yield. A total of 250 g of fresh plant parts material has been used. The essential oil obtained was dried over anhydrous sodium sulfate and kept at 4°C until analysis.

### Gas chromatography-mass spectrometry (GCFID/ GC-MS) analysis

The essential oil was analyzed by using GC-FID / MS analysis of the essential oils were done in Research Laboratory, Bingol University, by using HP (GC-MS) 7890A GC, 5975C MS and with a flame ionization detector FID detector were used. The study was performed by simultaneous injection in the same instrument. RXI-5 MS column (30m x 0,25 mm x 0,25 µm) Colon Flow Rate was used in both analyses with helium as the carrier gas in GC-MS. The injection volume was 1.0 μL of diluted solution (1 100^-1^) of oil in n-hexane. The temperature of the injector was 250°C, and the flow rate was 1.3 mL min^-1^. (splitless mode). The GC oven temperature was preserved at 70°C for 2 min and automated to 150°C at a rate of 10 °C/min and then kept constant at 150°C for 15 min to 240°C at a rate of 5°C min^-1^. A series of n-alkanes were used as reference points in the calculation of retention indices (RI). MS were occupied at 70 eV and a mass range of 35-425. The determination of the compounds was based on a comparison of their retention indices (RI), and mass spectra with those acquired from Wiley-Nist W9N11 libraries.

### Determination of the antimicrobial activity

In this study; *Staphylococcus aureus* ATCC25923, *Escherichia coli* ATCC25322, *Klebsiella pneumoniae* ATCC 700603, *Bacillus megaterium* DSM32, *Candida glabrata* ATCC 66032, *Candida albicans* FMC17, *Epidermophyton* sp., and *Trichophyton* sp. microorganisms were used. Microorganism cultures were obtained from Firat University, Faculty of Science, Department of Biology, Microbiology Laboratory culture collection. The antimicrobial activity of extracts plant obtained from methanol solvents was determined according to the disk diffusion method (Collins et al., 2004). *Bacteria strains* (*Escherichia coli* ATCC25922, *Klebsiella pneumoniae* ATCC700603, *Bacillus megaterium* DSM32, *Staphylococcus aureus* COWAN1, *Candida albicans* FMC17, *Candida glabrata* ATCC 66032, *Epidermophyton sp*. and *Trichophyton* sp) were incubated into Nutrient Broth (Difco) for 24 hours at 35 ± 1°C, yeast strains (*Candida albicans* FMC17 and *Candida glabrata* ATCC 66032) were incubated into Malt Extract Broth (Difco) and dermatophyte fungi (*Trichophyton* sp. and *Epidermophyton sp*.) was inoculated into Glucose Sabouroud Buyyon (Difco) and incubated at 25±1°C for 48 hours. The culture of prepared bacteria, yeast, and fungi in broth are respectively; was inoculated into Müeller Hinton Agar, Sabouraud Dextrose Agar, and Potato Dextrose Agar at a rate of 1% (10^6^ bacteria ml^-1^, 10^4^ yeast ml^-1^, 10^4^ fungi ml^-1^). Then, after shaking well, 25 ml was placed in sterile petri dishes of 9 cm diameter. Homogeneous distribution of the medium was achieved. 6 mm diameter antimicrobial discs (Oxoid), each impregnated with extracts of 100 µl (1000 µg) were lightly placed on the solidified agar medium. After the petri dishes prepared in this way were kept at 4°C for 1.5-2 hours, the plates inoculated with bacteria were incubated at 37 ± 0.1°C for 24 hours, and the plates were inoculated with yeast at 25 ± 0.1°C for 72 hours. As controls, different standard discs were used for bacteria (Streptomycin sulphate 10 µg disc^-1^) and yeasts (Nystatin 30 µg disc^-1^). Zones of inhibition were measured in mm.

### Determination of antioxidant activity

Total antioxidant status (TAS) and total oxidant status (TOS) of plant extracts were determined with Rel Assay kits (Rel Assay Kit Diagnostics, Turkey). TAS value was expressed as mmol Trolox equiv. L^-1^ and Trolox was used as the calibrator (Erel, 2004). The TOS value was expressed as μmol H_2_O_2_ equiv. L^-1^ and hydrogen peroxide was used as the calibrator (Erel, 2005). The antioxidant activity of the methanol extract of the plant was determined according to the 2.2-diphenyl-1-picrylhydrazyl (DPPH) radical scavenging capacity method (Cuendet et al., 1997; Kirby and Scmidt,1997). The solution was prepared in methanol at a concentration of 1 mg/ml of the extract obtained. The prepared solution was diluted four times and the calibration curve of DPPH was obtained. By taking 40 µl of the prepared solution, 160 µl of DPPH solution was added. After thorough mixing, the mouth was closed and kept in the dark for 30 minutes. The same procedures were repeated for all concentrations and methanol was used as a control. At the end of this period, the absorbances of each mixture were read at 570 nm in the spectrophotometer. % inhibition values were calculated;

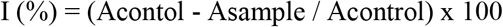

### Statistical analysis

Statistical analysis of the data was performed in the SPSS 22.0 package program and p<0.05 was regarded as statistically significant.

## RESULTS

### Essential oil studies

The composition of the aerial parts essential oils of *H. niger* and *H. niger* seeds are listed in Table 2, in which the percentage composition and retention index of components are given. *H. niger* seeds were hydrodistilled separately, obtaining yields of 0.5 % (v/w) of yellow oils. In the essential oils of *H. niger* seeds, 41 components were identified representing 99.0% of the oils. According to the results of the analysis hexahydrofarnesyl acetone 46.36%, hexanal 9.05 %as major components. The aerial part of *H. niger* was hydrodistilled, obtaining yields of 0.4% (v/w) of light yellowish oils. In the essential oils of the aerial part of *H. niger*, 22 components were identified representing 97.51% of the oils. More than half of the oil is phytol. According to the results of the analysis, phytol 52.09%, hexahydrofarnesyl acetone 19.66%, as major components. The analysis of the essential oil showed that sesquiterpenes in both oils were found. As a result of GS-MS, the essential oil components of the aerial parts and seeds of *H. niger* were determined in this study. These components are given in Table 2. The essential oils isolated were a complex mixture of sesquiterpenes and hydrocarbons. GC-MS was used to analyze the compositions of the volatile oil in the seeds.

**Table 2.**
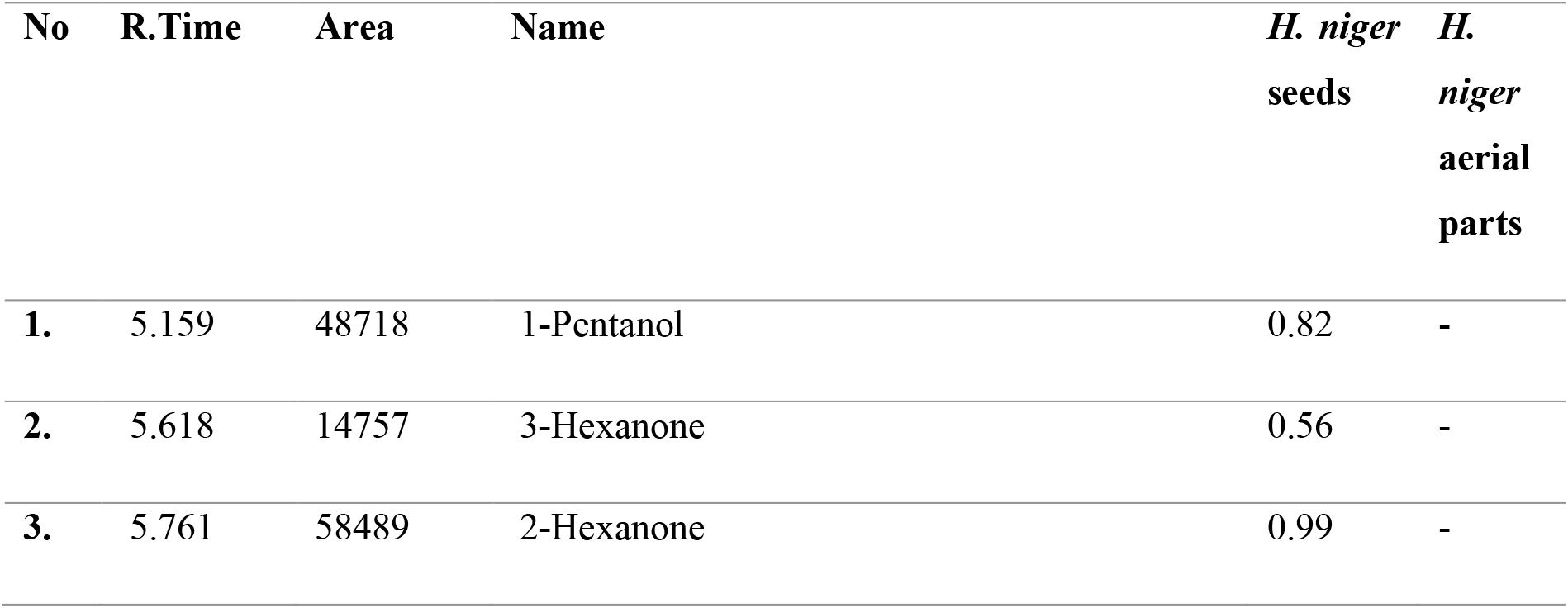
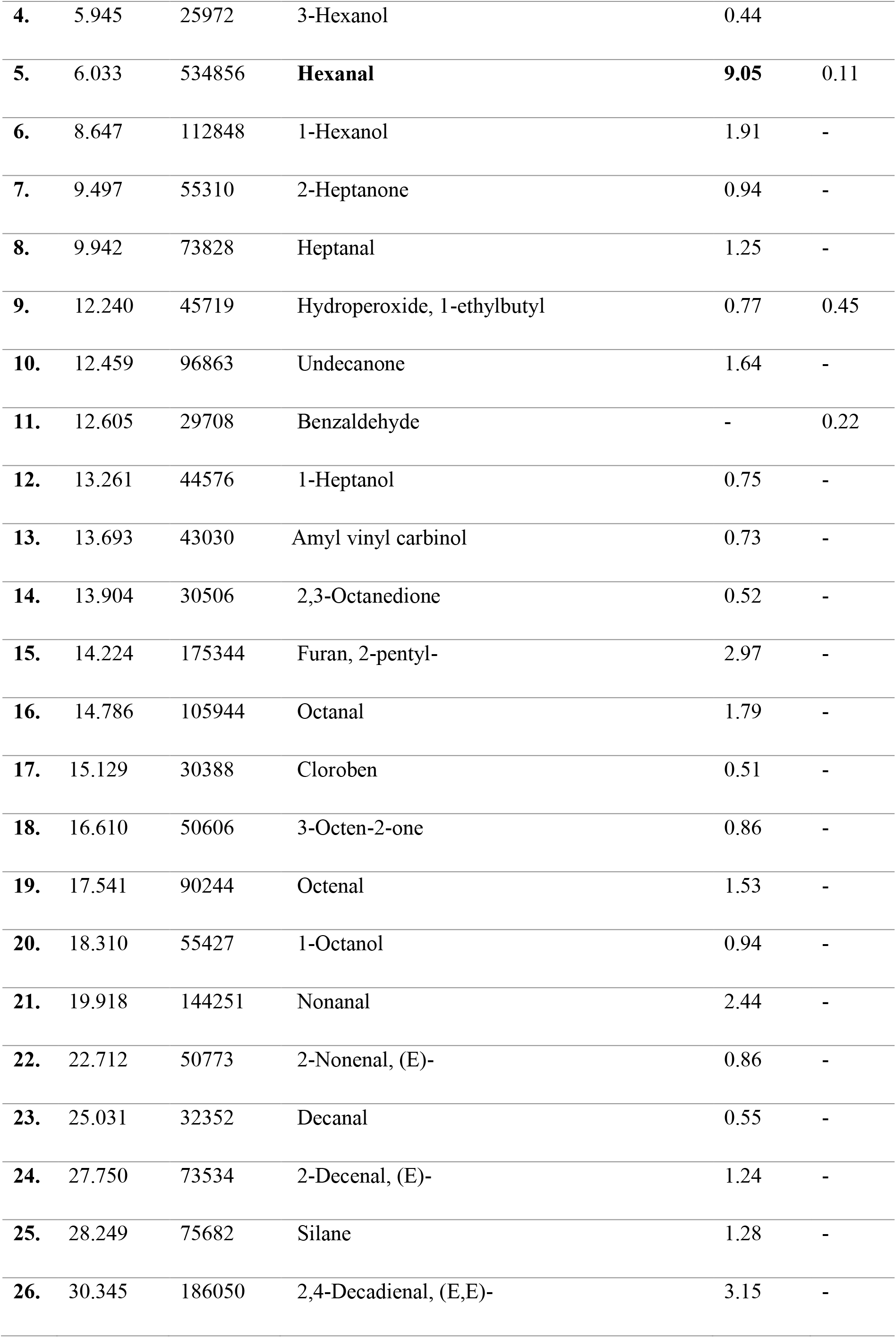
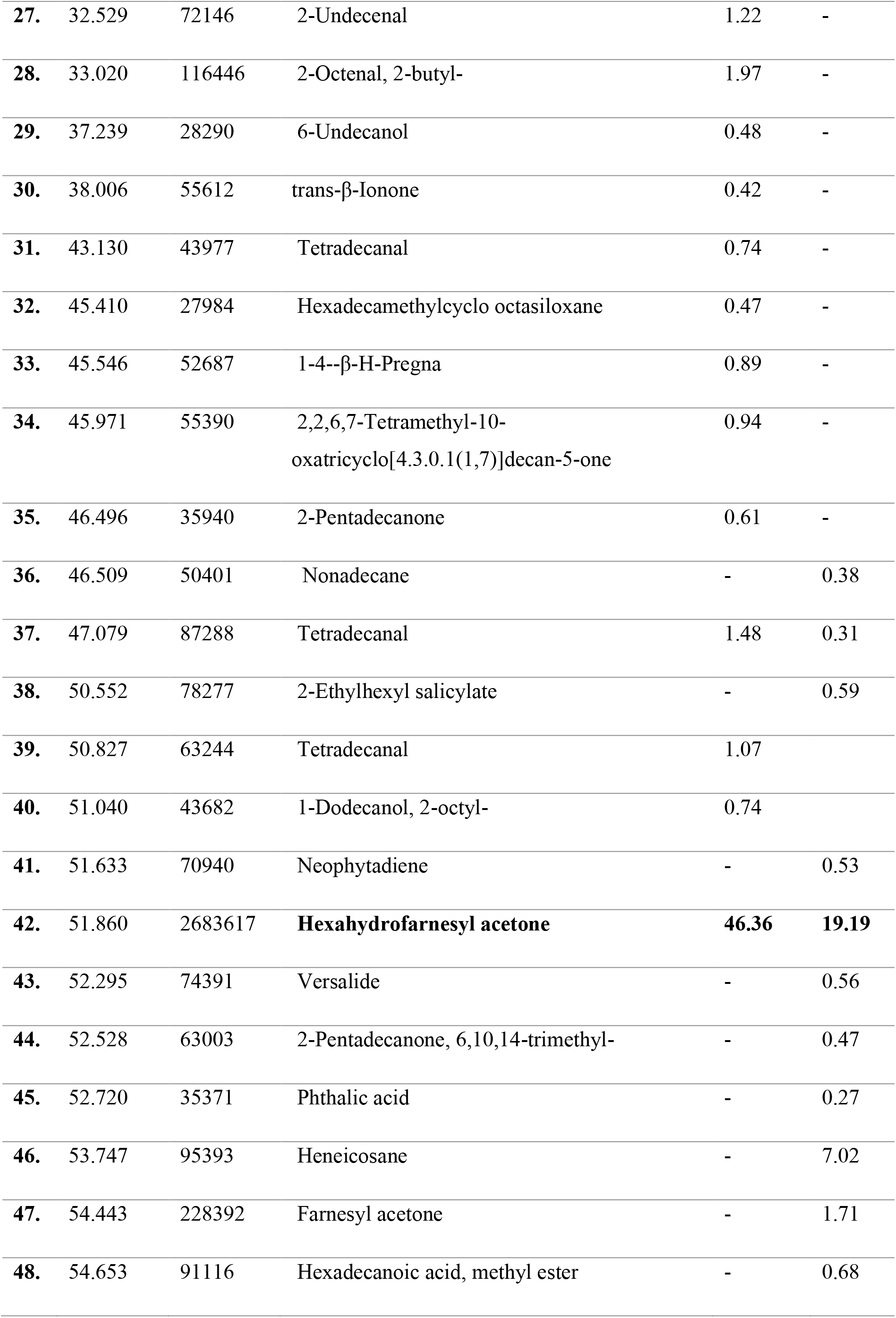
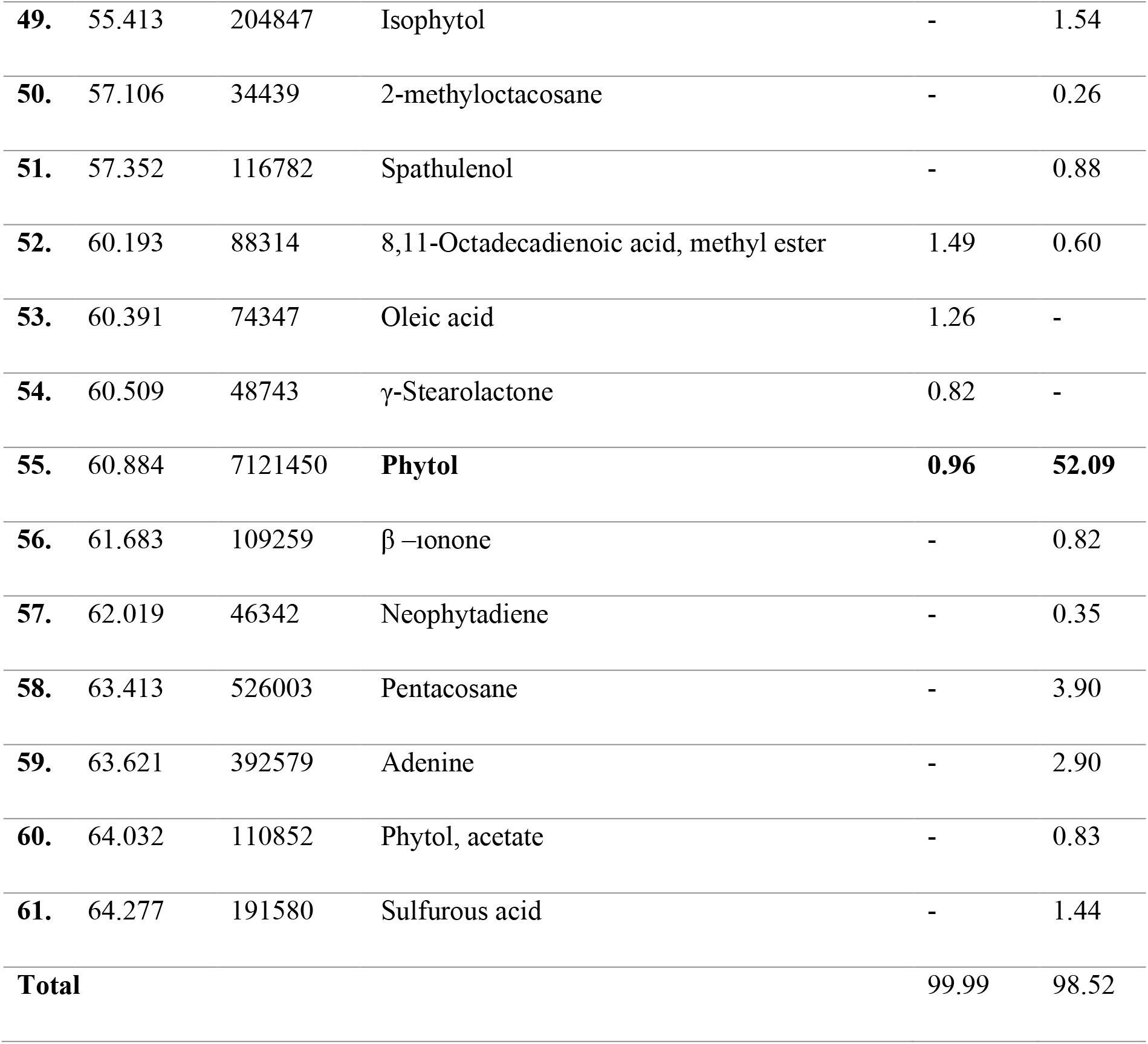
Constituents of the essential oils from *H. niger*

### Antimicrobial activity

The antimicrobial effect of *H. niger* against the microorganisms used is shown in Table 3. Methanol extract of *H. niger* created 32±0.11, 21±0.23, 20±0.51, 19±0.28, 23±0.17, 20±0.46, 13±0.46 and 16±0.23 mm inhibition zones against *E. coli, S. aureus, K. pneumoniae, B. megaterium, C. albicans, C. glabrata, Trichophyton sp*. and *Epidermophyton* sp., respectively. When the results obtained were compared with the controls, it was determined that it showed the best antimicrobial effect against *E. coli* (Table 3).

**Table 3.**
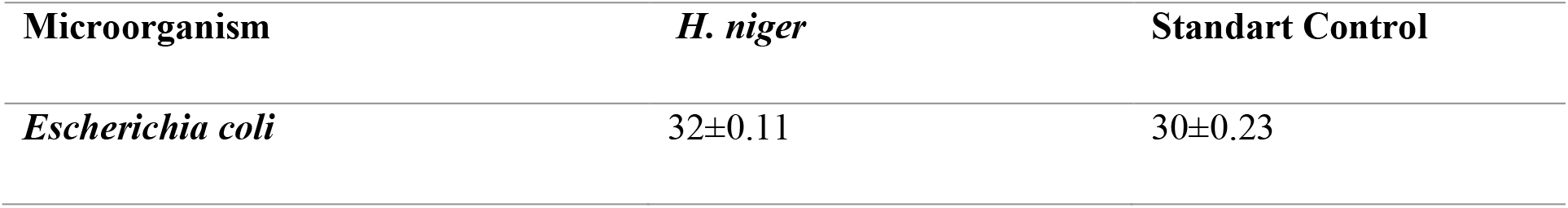
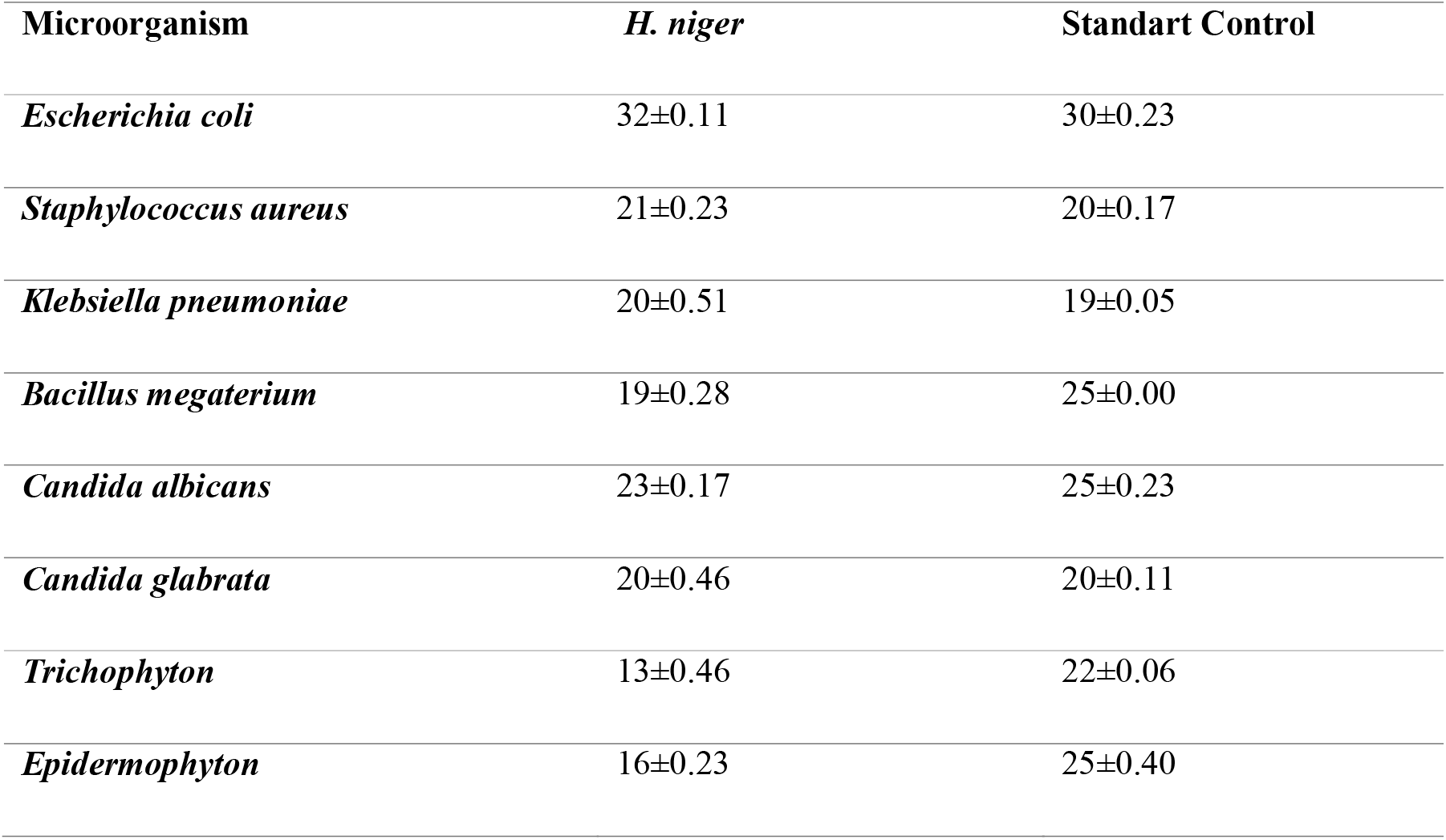
Antimicrobial effect of *H. niger*

### Antioxidant activity

While the TAS value of the methanol extract of *H. niger* at 1 mg ml^-1^ concentration was determined as 3.77±0.00 mmol, the TOS value was determined as 6.94±0.00 μmol at the same concentrations. In the results obtained, the TAS value of *H. niger* was very good, while the TOS value was found to be between normal values (Table 4). The scavenging effect of *H. niger* on DPPH radical is shown in Table 5. The DPPH radical scavenging effect of *H. niger*’s methanol extracts at 1000 µg mL^-1^ was determined as 73.07±0.02 %. 125 µg mL^-1^ showed a very low antioxidant effect. In the results obtained, it is seen that the scavenging effect of the DPPH radical increases depending on the increasing concentrations of *H. niger*.

**Table 4.**
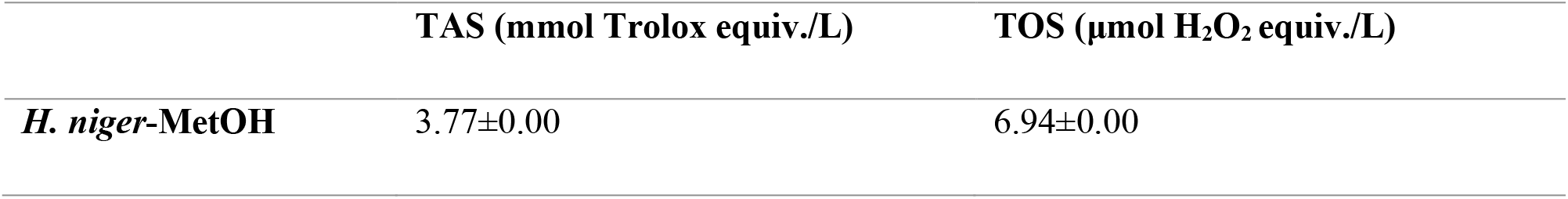
TAS and TOS values of *H. niger*

**Table 5.** Percent inhibition of the DPPH radical of root and leaf parts of *H. niger*

| Concentrations | *H. niger* -MetOH |
| --- | --- |
| 1000 µg/mL | 73.07±0.02 |
| 500 µg/mL | 52.73±0.07 |
| 250 µg/mL | 27.88±0.44 |
| 125 µg/mL | 15.78±0.29 |

## DISCUSSION

### Essential oil studies

*H. niger* leaves were studied in HPLC by extraction method, and 29 components were found and acid palmitic acid 28.30%, linoleic acid 26.85%, oleic acid 14.39 %, hexanal 10.24%, and stearic acid 1.81% were determined as major components (Wang et al., 2013). Compared with this study, our results show parallelism in terms of compounds that hexanal 9.05%, oleic acid 1.26%. However, the reason for the difference seen in other components (palmitic, linoleic acid) may be due to geographical and climatic differences. *H. muticus* L. subsp. *falezlez* aerial parts of essential oil analysis results, borneol (76.47%) was the main compound identified, followed by bornyl acetate 4.6% and 1,3-bis(2-cyclopropyl-2-methylcyclopropyl) but-2-en-1-one (4.19%) is the major substance. And they reported the oil yield was 0.50% (v w^-1^) in ten compounds (Ayari-Guentri et al., 2017). These substances were not found in our study. It can be said that the reason for this variability in the composition of essential oils is the differences in the geographical regions from which some species are taken and the variability of ecological conditions. These factors also play a role in the oil composition. In another study, the constituents of the most active methanol extracts of *H. niger* and *H. reticulatus* were determined by HPLC-Diode Array Detector-Electrospray Ionization Mass Spectroscopy. Three major phenolic constituents were identified and quantified: chlorogenic acid, rutin, and Quercetin-3o-Glucoside-Rhamnoside-Rhamnoside (Jassbi et al., 2014). The oil of *H. niger* and its methanol extract was showed antinematode activity (Kareem, 2020). The ethanol extract of *H. niger* seeds showed high insecticidal activity against the *Aphis laburni* and *Lucilia sericata* (Wang et al., 2006; Küpeli et al., 2020). Our results are compatible with the literature. So, secondary metabolites are generally synthesized in plants as they represent the most important defense mechanisms against pathogens; the amount produced along with the quality may vary as a function of habitat, the organ in which they are produced and climate conditions (Balogun et al., 2017; Moradi et al., 2020).

Hexahydrofarnesyl acetone is the major constituent of the *H. niger* seeds essential oil in this study. Hexahydrofarnesyl acetone has antibacterial, anti-nociceptive, and anti-inflammation activities (Wei, 2016). And the methanol extract of *H. niger* showed the highest antimicrobial effect. So, hexahydrofarnesyl acetone, which was a major compound present in all the oils extracted in this study, showed strong antimicrobial activity against gram-positive and gram-negative bacteria (Filipowicz et al., 2003); essential oil from *Equisetum arvense*, which was rich in hexahydrofarnesyl acetone, showed broad-spectrum inhibition against various bacterial and fungal species (Radulovic et al., 2006). Phytol, diterpene alcohol from chlorophyll, which is widely used as a food additive and in medical fields, has been found as a major substance in the aerial parts of *H. niger* essential oil in our study of phytol, which has promising antischistosomal properties. Phytol has antinociceptive and antioxidant activities as well as anti-inflammatory and antiallergic effects. And it showed that has an antimicrobial effect against *Mycobacterium tuberculosis* and *Staphylococcus aureus* (de Moras et al., 2014). As a result of our study, phytol, which is found at the rate of 52.09 percent in *H. niger* essential oil, was found to be high enough to form a source for other studies.

### Antimicrobial activity

The antimicrobial effect (26.2 mm-9.6 mm) of the seeds of *H. niger* against some microorganisms (*Enterococcus faecalis, Escherichia coli, Klebsiella pneumoniae, Pseudomonas aeruginosa, Proteus mirabilis*, and *Candida albicans*) was determined. The results obtained, showed the best antimicrobial effect against E. coli (Dulger and Dulger, 2015). In a different study, it was reported that *H. niger* has antifungal effects (Dulger et al., 2010). It was reported that the methanol extract obtained from the seeds of the same species showed strong antimicrobial activity (25.0 mm) against *S. aureus* (Dulger et al., 2010). It was reported that alkaloid extracts obtained from flower stems and roots of *H. niger* have strong antimicrobial effects (Chalabian et al., 2012). Crude protein extracts of the same species were tested against *E. coli, S. aureus, P. aeruginosa*, and *P. vulgaris*. It showed growth inhibition zone diameters of 14, 15, 14, and 20 mm against these pathogens, respectively (Mateen et al., 2015). It has been reported that *H. reticulatus* seed extract has an antimicrobial effect against *Bacillus subtilis, Escherichia coli, Klebsiella pneumoniae, Pasteurella multicoda, Pseudomonas aeruginosa, Salmonella enteritidis, Staphylococcus aureus, Yersinia enterocolitica* microorganisms (Akbaş et al., 2020). It has been determined that the leaf parts of give have a significant antimicrobial effect (Hossain et al. 2021). When the antimicrobial results of *H. niger* are compared with the literature studies, it is seen that the results obtained are different from each other. Alkaloids, glycosides, flavonoids, and their derivatives, which are usually found in plant extracts, are responsible for the antimicrobial effect (Dulger et al., 2010a-b; Benhouda et al., 2014; Dulger and Dulger, 2015; Ayari-Guentri et al., 2017; Elsharkawy et al., 2018; Akbaş et al., 2020). In addition, the plant species, concentrations, and microorganisms used to affect the antimicrobial results.

### Antioxidant activity

The DPPH radical scavenging effect of the ethanol extract of *H. niger* was reported as 2% at a concentration of 10 mg/ml however, this value gradually decreases at high concentrations (Hajipoor *et al*., 2015). The antioxidant activity of the aerial parts of *Hyoscyamus niger* extracts was investigated by DPPH (2, 2-diphenyl-1-picrylhydrazyl) and iron-reducing antioxidant power (FRAP) activities. Antioxidant (EC50) for methanol extract was 377±1.21 μg ml^-1^, 21±0.68, and was 4.8±0.32 μg ml^-1^ for butylated hydroxytoluene and ascorbic acid (Koçpınar, 2020). Free radical scavenging activity of seven fractions of *Hyoscyamus niger* alkaloidal extract was investigated by 2, 2-diphenyl-1-picrylhydrazyl activity. Only a fraction of the alkaloidal extract showed moderate free radical scavenging activity compared to positive and negative controls (Jassbi et al., 2014). Methanolic extracts of *Hyoscyamus niger* showed antioxidant activity (IC50=1.64 μg) compared to α-tocopherol (IC50=0.60 μg) used as positive control (Souri et al. 2004). The IC50 values of the DPPH radical scavenging effect of the methanol extracts of the seed, leaf, and epicalyx parts of *H. niger* were determined as 1485±34.9, 270.2±8.3, and 451.3±2.2 μg mL^-1^, respectively (Hajipoor et al., 2015). The methanolic extract of *Hyoscyamus albus* leaves displayed the highest antioxidant activity (76.00%) (Benhouda et al., 2014). *Hyoscyamus reticulatus* aqueous extract exhibited significant antioxidant scavenging properties (533.26 μmol TE g^-1^ dry extract weight) (Akbaş et al., 2020). The antioxidant testing showed that the methanolic extract of *H. muticus* has a noticeable antioxidant activity with an IC50 of 8.1±0.65 mg/ml and an EC50 of 12.74±1.12 mg ml^-1^ (Dehghan et al., 2012) The results of DPPH radical scavenging activity of *H. muticus* L. subsp. *falezlez* methanolic extracts showed the highest IC50 value of stem (0.54 ± 0.19 mg mL^-1^), followed by leaf (0.65 ± 0.29 mg mL^-1^) and flower (0.92 ± 0.39 mg mL^-1^) (Ayari-Guentri et al., 2017). When the results obtained are compared with the literature, it is seen that the antioxidant effects vary according to the type used. Because of the plant species used, the habitat collected and the presence of bioactive components in the plant materials affect the antioxidant results.

## CONCLUSIONS

It was determined that the highest antimicrobial effect of *H. niger* was against *E. coli*. When the study results are compared with previous studies, the results vary depending on the parts of the plant material used, the microorganism, the solvent, and the concentrations. It was determined that the antioxidant effect of the species is good at 1 mg ml^-1^ concentration. When the results are compared with the literature, it is seen that there are some differences. Because it varies depending on the parts of the plant materials used, the concentration, and the solvent. *Hyoscyamus* species are known to have important pharmacological effects thanks to the phytochemicals they contain. It is important to investigate these at a sufficient level and especially to evaluate the phytochemicals they contain in terms of bringing them to the literature. It is a medicinal plant that should be studied for this purpose in *H. niger*. The bioactive substances, especially hexahydrofarnesyl acetone and phytol, in the essential oils from seeds and aerial part *H. niger* may be of therapeutic interest.

## Competing Intererest

The authors declare no competing or financial interests.

## Author contributions

Şİ, PYS, AD, SK SC, designed the study and experiments and analyzed the data conceived and and drafted and revised the paper.

## References

Akbaş, P., Uslu, H., Uslu, G. A. and Alkan, H. (2020). Hyoscyamus reticulatus L. tohum ekstraktının antimikrobiyal ve apoptotik etkinliğinin araştırılması. Süleyman Demirel Uni. J. Nat. Appl. Sci. 24, 317–322. doi:10.19113/sdufenbed.631074

Ayari-Guentri, S., Djemouai, N., Gaceb-Terrak, R. and Rahmania, F. (2017). Chemical composition and antioxidant activity of Hyoscyamus muticus L. subsp. falezlez (Coss.) Maire from Algeria. J. Essential Oil Bearing Plants. 20, 1370–1379. doi:10.1080/0972060X.2017.1396930.

Balogun, O. S., Solomon Ajayi, O. and Adeleke, A. J. (2017). Hexahydrofarnesyl acetone-rich extractives from Hildegardia barteri. J. Herbs, Spıces & Med Plants. 23, 393–400. doi:10.1080/10496475.2017.1350614.

Chalabian, F., Majd, A., Mehrabian, S. and Falahian, F. (2002). A study of growth inhibitory effect of alkaloids of two species of genus Hyoscyamus on some kinds of microbes of skin. J. Scı. 12, 3371–3378.

Collins, C. H., Lyne, P. M., Grange, J. M. and Flkinham III, J. O. (2004). Microbiological methods, eight ed, London.

Cuendet, M., Hostettmann, K., Potterat, O. and Dyatmiko, W. (1997). Iridoid glucosides with free radical scavenging properties from Fagraea blumei. Helvetica Chim. Acta. 80, 1144–1152. doi:10.1002/hlca.19970800411

de Moraes, J., de Oliveira, R. N., Costa, J. P., Junior, A. L., de Sousa, D. P., Freitas, R. M., Allegretti, S. M. and Pinto, P. L. S. (2014). Phytol, a diterpene alcohol from chlorophyll, as a drug against neglected tropical diseaseSchistosomiasis mansoni. PLoS Negl. Trop. Dis. 8, e2617. doi:10.1371/journal.pntd.0002617

Dehghan, E., Häkkinen, S. T., Oksman-Caldentey, K. M. and Ahmadi, F. S. (2012). Production of tropane alkaloids in diploid and tetraploid plants and in vitro hairy root cultures of Egyptian henbane (Hyoscyamus muticus L.). PCTOC. 110, 35–44.

Demirpolat, A. B., Aydoğmuş, E. and Arslanoğlu, H. (2021). Drying behavior for Ocimum basilicum Lamiaceae with the new system: exergy analysis and RSM modeling. Biomass Conversion and Biorefinery. 12, 515–526. doi:10.1007/s13399-021-02010.

Dulger, B., Goncu, B. S. and Gucin, F. (2010a). Antibacterial activity of the seeds of Hyoscyamus niger L. (Henbane). Asian J. Chem. 22, 6879–6883.

Dulger, B., Hacioglu, N., Goncu, B. S. and Gucin, F. (2010b). Antifungal activity of seeds of Hyoscyamus niger L. (Henbane) against some clinically relevant fungal pathogens. Asian J. Chem. 22, 6321–6324.

Dulger, G. and Dulger, B. (2015). Antimicrobial activity of the seeds of Hyoscyamus niger L. (Henbane) on microorganisms isolated from urinary tract infections. J. Med. Plants Studies. 3, 92–95.

Erel, O. (2004). A novel automated direct measurement method for total antioxidant capacity using a new generation, more stable ABTS radical cation. Clin. Biochem. 37, 277–285. doi:10.1016/j.clinbiochem.2003.11.015

Erel, O. (2005). A new automated colorimetric method for measuring total oxidant status. Clin. Biochem. 38, 1103–1111. doi:10.1016/j.clinbiochem.2005.08.008

Filipowicz, N., Kamiński, M., Kurlenda, J., Asztemborska, M. and Ochocka, J. R. (2003). Antibacterial and antifungal activity of juniper berry oil and its selected components. Phytother Res. 17, 227–231. doi:10.1002/ptr.1110

Guha, S. and Maheshwari, S. C. (1966). Cell division and differentiation of embryos in the pollen grains of Datura in vitro. Nature. 212, 97–98

Haas, L. F. (1995). Hyoscyamus niger (henbane). J. Neurol. Neurosurg. Psychiatry. 59, 114. doi:10.1136/jnnp.59.2.114

Hajipoor, K., Sani, A. M. and Mohammad, A. (2015). In vitro antioxidant activity and phenolic profile of Hyoscyamus niger. IJBPAS. 4, 4882–4890.

Hossain, M. A. and Al-Touby, S. S. J. (2021). A phytopharmacological review on an important indigenous medicinal plant Hyoscyamus gallagheri. Arabian J. Med. and Aromatic Plants. 7, 367–384. doi:10.48347/IMIST.PRSM/ajmap-v7i3.26721

Ismeel, A. O. (2011). Cytogenetic and cytotoxic studies on the effect of phytoinvestigated active compounds of Hyoscyamus niger (in vivo and ex vivo). Al-Nahrain University-College of Science, Iraq.

Jassbi, A. R., Miri, R., Masroorbabanari, M., Asadollahi, M., Attarroshan, M. and Baldwin, I. T. (2014). HPLC-DAD-ESIMS Analyses of Hyoscyamus niger and H. reticulatus for their antioxidant constituents. Austin Chromatogr. 1, 1022.

John, H., Binder, T., Höchstetter, H. and Thiermann, H. (2010). LC-ESI MS/MS quantification of atropine and six other antimuscarinic tropane alkaloids in plasma. Analytical and Bioanalytical Chem. 396, 751–763.

Kareem, Z. J. (2020). Biomedical applications and secondary metabolite profiling of Hyoscyamus niger and Sesamum indicum seed, root and hairy root cultures. Georg-August-Universität Göttingen.

Kirby, A. J. and Schmidt, R. J. (1997). The antioxidant activity of Chinese herbs for eczema and of placebo herbs. I. J. Ethnopharma. 56, 103–8. doi:10.1016/S0378-8741(97)01510-9

Koçpınar, E. F. (2020). Antioxidant behavior of Hyoscyamus niger having narcotic effect on heavy metal reduction and radical scavenging. J. Physical Chem. and Functional Materials. 3, 68–71.

Kosari, M., Noureddini, M., Khamechi, S. P., Najafi, A., Ghaderi, A., Sehat, M. and Banafshe, H. R. (2021). The effect of propolis plus Hyoscyamus niger L. methanolic extract on clinical symptoms in patients with acute respiratory syndrome suspected to COVID-19: A clinical trial. Phytother. Res. 35, 4000–4006. doi:10.1002/ptr.7116

Küpeli Akkol, E., Ilhan, M., Kozan, E., Gürağaç Dereli, F. T., Sak, M. and Sobarzo-Sánchez, E. (2020). Insecticidal Activity of Hyoscyamus niger L. on Lucilia sericata Causing Myiasis. Plants. 9, 655. doi:10.3390/plants9050655

Li, J., Shi, J., Yu, X., Sun, J., Men, Q. and Kang, T. (2011). Chemical and pharmacological researches on Hyoscyamus niger. Chinese Herbal Med. 2011, 117–126.

Maiti, R. K., Cerda, V., Villarreal, R. and Treviño, V. (2002). Some aspects on pharmacognosy of ten species of the family solanaceae utiliz. Caldasia. 317-321.

Mateen, A., Tanveer, Z., Janardhan, K. and Gupta, V. C. (2015). Screening and purification of antibacterial proteins and peptides from some of the medicinal plants seeds. Int. J. Pharma and Bio. Scie. 6, 774–781.

Moradi, H., Ghavam, M. and Tavili, A. (2020). A. Study of antioxidant activity and some herbal compounds of Dracocephalum kotschyi Boiss. in different ages of growth. Biotechnol. Reports. 25, e00408. doi:10.1016/j.btre.2019.e00408

Orbak, Z., Tan, H., Karakelleoğlu, C., Alp, H. and Akdag R. (1998). Hyoscyamus niger (henbane) poisonings in the rural area of east Turkey. Headache. 14, 6–5.

Pudersell, K., Vardja, T., Vardja, R., Matto, V., Arak, E. and Raal, A. (2012). Inorganic ions in the medium modify tropane alkaloids and riboflavin output in Hyoscyamus niger root cultures. Pharmacognosy Magazine. 8, 73. doi:10.4103/0973-1296.93330

Radulović, N., Stojanović, G. and Palić, R. (2006). Composition and antimicrobial activity of Equisetum arvense L. essential oil. Phytother. Res. 20, 85–88. doi:10.1002/ptr.1815

Souri, E., Amin, G., Dehmobed-Sharifabadi, A., Nazifi, A. and Farsam, H. (2004). Antioxidative activity of sixty plants from Iran. Iranian J. Pharma Res. 3, 55–59. doi:10.22037/IJPR.2010.298

Tanker, N., Koyuncu, M. and Coşkun, M. (1998). Farmasötik botanik. Ankara Üniviversitesi Eczacılık Fakültesi Yayınları, Ankara.

Ugur, T. (2013). Tıbbi olarak kullanılan bazı bitkilerin kanser hücreleri üzerinde sitotoksik etkilerinin araştırılması. Muğla.

Wang, X., Wang, Y., Li, J., Men, Q. and Kang, T. (2013). Analysis of volatile oil in processed samples of seeds of Hyoscyamus niger with GC-MS. Chinese Archives of Traditional Chinese Med. 05.

Wang, Z. J., Zhou, J., Li, W. J., Yan, Y. J., Dai, B. and Fan, Z. J. (2006). Insecticidal activity of extracts from plant recourses in Xinjiang. Modern Agrochemicals. 5, 34.

Wei, G., Kong, L., Zhang, J., Ma, C., Wu, X., Li, X. and Jiang, H. (2016). Essential oil composition and antibacterial activity of Lindera nacusua (D. Don) Merr. Nat. Product. Res. 30, 2704–2706. doi:10.1080/14786419.2015.1135145

Yücel, U. M. and Yılmaz, O. (2014). Total alkaloid amounts of two Hyoscyamus species (Henbane) grown in Van region. Van Veterinary J. 25, 77–80.

